# Localisation and hypersecretion of Nerve Growth Factor in breast Phyllodes tumours: evidence from a preliminary study

**DOI:** 10.1101/392100

**Authors:** Ashutosh Kumar, Khursheed Raza, Tapas C. Nag, Anurag Srivastava, Ritu Sehgal

## Abstract

**Background:** The pathophysiology of the breast phyllodes tumours is uncertain. Currently, wide surgical removal is the only available treatment option. The histopathological diagnosis of phyllodes tumours is often confused with that of fibroadenomas due to a striking histological resemblance; hence a distinctive biomarker for this tumour type is warranted.

**Material & Methods:** Fresh human breast tissue was obtained from surgically excised breast phyllodes and fibroadenoma tumours (test, 2 cases each), breast cancer (positive control, 2 cases) and normal breast tissue (negative control, 1 case). Immunohistochemistry was performed for the detection of nerve growth factor (NGF) on frozen sections of the test and control tissues fixed in 4% paraformaldehyde, using the indirect streptavidin-biotin-peroxidase complex method. Sandwich ELISA on tissue homogenates of the same test and control cases was also performed to validate the immunohistochemical findings.

**Results:** A marked difference in NGF expression was detected in phyllodes tumours compared to fibroadenomas. The maximum NGF expression was observed in phyllodes tissue followed by cancer tissue, and the least expression in fibroadenomas (3-5 times less than in phyllodes; comparable with normal breast tissue).

**Conclusion:** NGF is known for its growth inducing potential in breast cancer, but its secretion by a benign breast tumour is not known in literature. This study reports abundant NGF secretion by breast phyllodes, raising the possibility of its potential role in tumour pathogenesis and progression that can be exploited therapeutically in future. We also propose that NGF may be used as a distinct biomarker of phyllodes tumours, for differentiating them from fibroadenomas during histopathology.

## Introduction

Phyllodes are rare fibro-epithelial tumours of the breast with an incidence of less than 1% of which malignant phyllodes account for 10-20% of the tumours [1]. The phyllodes tumours show high potential for growth, recurrence and malignant transformation, being sub-classified into three categories: benign, borderline and malignant [2]. Lymph node involvement and distant metastasis are rare but known in malignant cases.

Knowledge about phyllodes tumorigenesis is quite patchy and scarce in available literature. No biomarker or drug target has yet been identified for these tumours. This article reports for the first time, new evidence of hypersecretion of nerve growth factor (NGF) by phyllodes tumours. NGF is known to be secreted exclusively by cancer cells (including breast cancer), and suspected to be involved in their etiogenesis and tumour progression [3, 4]. Excessive secretion by benign breast phyllodes as observed in our study implies NGF involvement in the genesis and progression of these tumours. We also propose a potential diagnostic value for NGF as a biomarker to differentiate phyllodes tumours from fibroadenomas. These two benign growths share a very close histological resemblance that leads to frequent errors in histopathological diagnosis [5, 6].

## Material and methods

### Material

The test samples of benign phyllodes and fibroadenoma (2 cases each) were obtained from fresh surgical specimens of breast tumours removed from clinically and histopathologically diagnosed cases in women. The positive controls (2 cases) and the single negative control were taken from established cases of malignant (intra-ductal carcinoma) and normal breast tissue, respectively. The controls were chosen based on their known NGF secretory status, as reported in the literature [3, 4]. All these samples were provided by the Departments of Surgery and Pathology, All India Institute of Medical Sciences (AIIMS), New Delhi after ethical clearance from the institutional human ethics committee (IEC No: NP-325/2013RP03/2013).

### Methods

#### Immunohistochemistry (IHC)

IHC was performed on cryosections (thickness: 12-14µm) of tissue samples fixed in 4% paraformaldehyde in 0.1 M phosphate buffer for 6-8 hours at 4°C, using the standard indirect streptavidin-biotin-peroxidase complex method. The sections were incubated with primary antibodies against NGF (dilution: 1 µg/ml, Abcam Plc., UK, rabbit polyclonal) for 48 h, followed by incubation in anti-rabbit secondary antibody (dilution: 1 µg/ml) and avidin-biotin peroxidase (Vectastain Elite ABC kit, Vector Laboratories, Burlingame, CA, USA). The reactions were developed by treating sections with 3, 3 diaminobenzidine tetrahydrochloride (DAB, Sigma-Aldrich Corporation, MO, USA). The core biopsy tissue of the breast cancer and normal breast tissue were used as external positive and negative controls, respectively. The internal negative control was the tumour tissue without applying primary antibody. Haematoxylin was used as a counter-stain in fibroadenoma cases to see histological details. The images were captured with a digital camera under an optical microscope (Leica DM6000B bright-field microscope) using software (Leica Application Suites, Version 3.4.1; Leica Microsystems, Switzerland). The images were analysed using ImageJ software (NIH, USA) for assessment of staining intensity in specific tissue components and overall scoring.

#### Enzyme-linked immunosorbent assay (ELISA)

ELISA (human beta NGF, sensitivity <14 pg/ml, range 6.86 pg/ml to 5000 pg/ml; catalog number: ab99986, Abcam Plc., UK) was applied on test and control samples for quantification of NGF. Fresh surgically removed tissue samples were preserved with isopentane at − 80 ºC. The tissue were thawed, weighed and homogenised in homogenization buffer (10 mM Tris, 150 mM NaCl, 0.25% sodium deoxycholate and protease inhibitors). Homogenates were centrifuged at 10,000 rpm at 4^°^C, following which the supernatant was collected for total protein estimation by Bradford method and equilibration using assay buffer. NGF competitive inhibition ELISA was performed in duplicates using the antibody pre-coated plate, by following the protocol provided with the kit. The readings of the test were taken at 450 nm in an ELISA plate reader and the data analysed for final NGF concentration in the samples, using the standard curve values derived from the recombinant NGF provided with the kit (concentration was expressed in ng/50µl).

### Results

#### Immunohistochemistry (IHC)

Intense NGF expression was observed in test samples obtained from cases of benign phyllodes, comparable with that seen in breast tissue collected from cases of intra-ductal carcinoma (IDC, positive control). However, NGF expression was almost absent in test samples obtained from cases of fibroadenoma (Fig. 1, Table 1).

**Table 1:** NGF expression in breast tumours (as detected by IHC)

| Tissue components | CA Breast | Phyllodes | Fibroadenoma | Normal breast tissue |
| --- | --- | --- | --- | --- |
| Duct | +++ | +++ | — | — |
| Stroma | ++ | +++ | — | — |
| Adipocytes | ++ | +++ | — | — |
| Small vessels | +++ | +++ | — | — |
| ***Total score*** | 10 | 12 | 0 | 0 |

— (No expression detected, score = 0), + (weak expression, score = 1), ++ (moderate expression, score = 2), +++ (intense expression, score = 3), CA (carcinoma), IHC (Immunohistochemistry)

**Figure 1:**
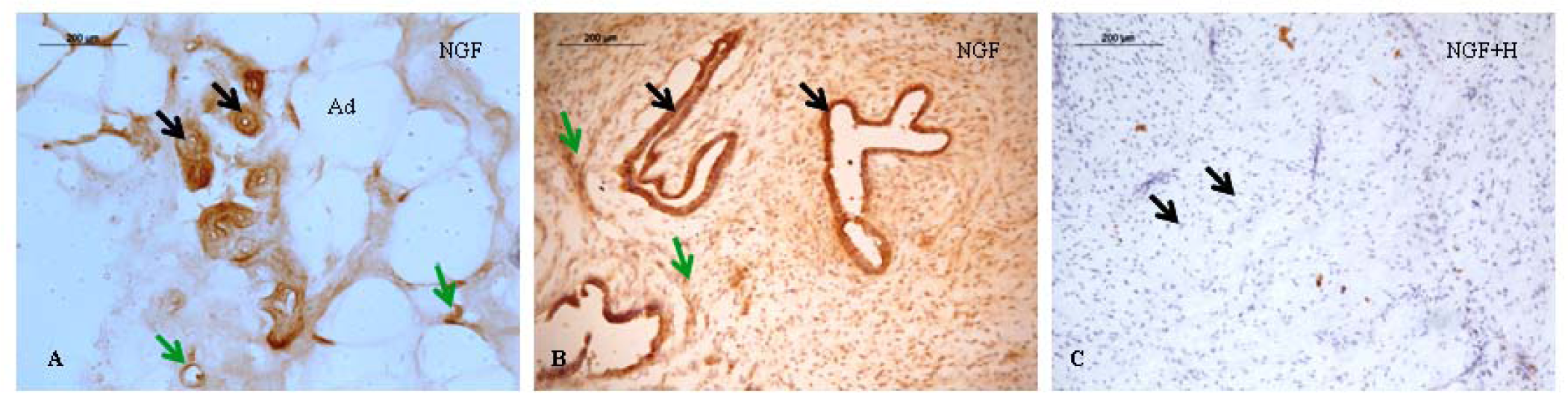
Immunohistochemical localisation of NGF in breast tumours. **A.** CA breast (IDC), intense NGF expression in ductal epithelium (black arrows) and small vessels (green arrows), Ad – adipocyte. **B.** Phyllodes tumour, intense expression in ductal epithelium (black arrows) and stroma (green arrows). **C.** Fibroadenoma, negligible expression, tissue counterstained with hematoxylin (H) to show nuclei of fibrocytes, characteristic of fibroadenoma (black arrows).

#### Enzyme-linked immunosorbent assay (ELISA)

Quantification of NGF secretion gave readings that matched with IHC scores for all the test and control samples. The highest NGF levels were observed in phyllodes tumours, followed by breast carcinoma (CA). The levels were low and quite comparable in tissue specimens obtained from fibroadenomas and normal breast (Fig. 2).

**Figure 2:**
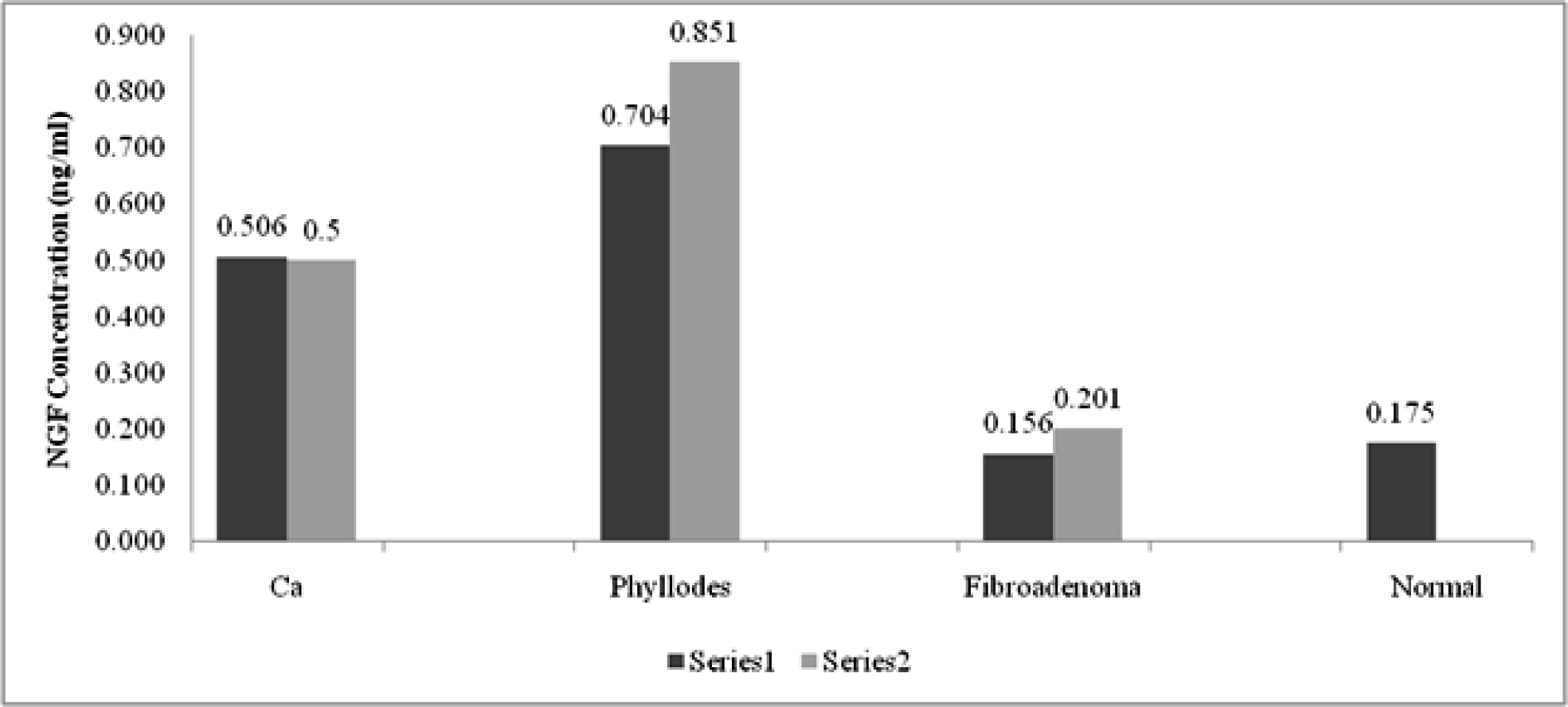
NGF expression in breast tumours (as detected by ELISA) Ca – breast cancer, Series 1 & 2 refers to the tissue samples collected from separate cases.

## Discussion

NGF is known for its growth-inducing potential and has been implicated for tumour progression in CA breast [4]. Although NGF has not been directly implicated in the pathogenesis of phyllodes tumours of the breast so far, a thorough literature survey revealed that certain molecules and pathways reported for these tumours can be structurally or functionally linked with NGF signalling. We observed a generalised over-expression on IHC in all tissue components of phyllodes tumours (ductal layers, stromal components, vessels) and negligible expression in those of fibroadenoma (Fig. 1, Table 1). Quantittive estimation by ELISA revealed NGF levels in phyllodes tumours to be approximately 1.5 times those observed in CA breast and 3-5 times those seen in fibroadenoma. This singular NGF over-expression by phyllodes tumours of the breast that has been noted in this study, points towards its possible role in tumourigenesis and tumour progression, and suggests the use of this molecule as a potential drug target. Jardim *et al.* reported activating mutations for N-RAS oncogene and intense expression of Protein Kinase B (Akt) and mammalian target of rapamycin (mTOR) in malignant phyllodes with a concomitant over-expression of phosphoinositide 3-kinase (PI3K) [7]. Tyrosine kinase A (TrkA) is a high affinity receptor for NGF that mediates growth and proliferation. N-RAS is an oncogene which is thought to be an essential molecule for executing NGF-TrkA mediated PI3K/Akt/m-TOR activation and NGF mediated differentiation of PC12 cells (from a neuron-like cell line derived from pheochromocytoma of the rat adrenal medulla) [8]. Coincidently, PI3K/Akt/m-TOR is the chief pathway mediating NGF signalling involved in growth and proliferation [9]. Furthermore, N-RAS has the chromosomal location 1p13.2 (Gene ID: 4893, NCBI, 2016) very close to that of NGF which is 1p13.1 (Gene ID: 4803, NCBI, 2016), suggesting the possibility of a functional linkage between the two genes [10]. N-RAS mutations are common in various tumours and a point mutation of the N-RAS gene may cause constitutive activation of the N-RAS dependent signalling pathways [11].

Several researchers have also found a huge genomic instability in phyllodes tumours which includes loci of 1p and 1q among other chromosomes [7, 12]. TrkA, the high affinity NGF receptor that mediates growth and proliferation is located on chromosome 1q 21-22. Moreover, keeping in mind the above-mentioned proximity of gene loci for NGF (1p13.1) and N-RAS (1p13.2), the instability of either of these loci may influence NGF expression and signalling in phyllodes tumours. Hence the role of this proximity of loci and concomitant genomic instability in the pathogenesis of phyllodes tumours needs to be examined.

### Differentiation of phyllodes from fibroadenomas

Phyllodes tumours are difficult to differentiate histologically from fibroadenomas [2, 6]. Although pronounced stromal cell activation is characteristic of phyllodes, sometimes it becomes truly puzzling for the pathologist to make an accurate diagnosis in cases where the tumours have intermediate features [5, 6]. The unambiguous NGF over-expression by phyllodes (3-5 times more than that by fibroadenoma) that was observed in the present study, makes NGF a highly likely candidate for a suitable biomarker of breast phyllodes.

### Limitations and further considerations

The sample size of this study being small, further studies with adequate sample size are necessary to validate the results. Future research may elucidate the dynamic interaction of NGF signalling pathway molecules in breast phyllodes. It would be particularly relevant and rewarding to establish beyond doubt, the diagnostic efficacy and specificity of NGF as a potential biomarker for phyllodes tumours of the breast.

## Abbreviations

NGF: nerve growth factor
TrkA: tyrosine kinase A receptor
IHC: immunohistochemistry
ELISA: enzyme-linked immunosorbent assay
IDC: intra-ductal carcinoma
CA: carcinoma
N-RAS: oncogene for N-ras protein that regulates cell division
Akt: Ak (mouse strain) transforming (retrovirus)/Protein Kinase B
m-TOR: mammalian target of rapamycin
PI3K: phosphoinositide 3-kinase
p: short arm of chromosome
q: long arm of chromosome
PC12: neuron-like cell line derived from pheochromocytoma of the rat adrenal medulla
NCBI: National Center for Biotechnology Information

## Declarations

### Ethics approval and consent to participate

Ethics approval from the Institute Ethics Committee (IEC) of the All India Institute of Medical Sciences (AIIMS) and informed consent from the participants was duly acquired prior to the commencement of the study (IEC No: NP-325/2013RP03/2013, AIIMS, New Delhi, India).

### Consent for publication

Not applicable.

### Availability of data and material

The datasets used and analyzed during the current study may be acquired from the corresponding author on reasonable request.

### Authors’ contributions

AK designed the study, performed the experiments, analyzed the data and wrote the manuscript. KR assisted in the experiments, data analysis and manuscript preparation. TCN provided the benefit of expertise and practical assistance in setting up the experiments and acquiring images. TCN, AS and RS edited the manuscript and provided logistic support and requisite guidance for designing and conducting the study. All authors read and approved the final manuscript.

## Acknowledgements

We sincerely thank Professor T. S. Roy and Dr. T. G. Jacob from the Department of Anatomy, AIIMS, New Delhi. Dr. Roy gave unstinted support and practical assistance in helping us access the resources necessary for conducting this study. Dr. Jacob provided valuable inputs for data analysis and manuscript writing.

